# A Chimeric Peptide Inhibits Red Blood Cell Invasion by Plasmodium falciparum with Hundredfold Increased Efficacy

**DOI:** 10.1101/2021.09.28.462119

**Authors:** Jamsad Mannuthodikayil, Suman Sinha, Sameer Singh, Anamika Biswas, Irshad Ali, Purna Chandra Mashurabad, Wahida Tabassum, Pratap Vydyam, Mrinal Kanti Bhattacharyya, Kalyaneswar Mandal

## Abstract

Inhibition of tight junction formation between two malaria parasite proteins, apical membrane antigen 1 and rhoptry neck protein 2, crucial for red blood cell invasion, prevents the disease progression. In this work, we have utilized a unique approach to design a chimeric peptide, prepared by fusion of the best features of two peptide inhibitors, that has displayed parasite growth inhibition, *in-vitro*, with nanomolar IC_50_, which is hundredfold better than any of its parent peptides. Further, to gain structural insights, we computationally modeled the hybrid peptide on its receptor.

---

Malaria infects over 200 million people and leads to nearly half a million death every year.^1^ Widespread mutations and deletions in the *Plasmodium* genome adversely affect the treatment options for the deadly disease. Among four major *Plasmodium* species that cause malaria in human, the *Plasmodium falciparum* (*Pf*) accounts for most malaria-related deaths worldwide. The recent emergence of several drug-resistant *Pf* strains makes this parasite the greatest threat to global health. The tight junction formation between two parasite proteins, apical membrane antigen 1 (*Pf*AMA1) and rhoptry neck protein 2 (*Pf*RON2), is known to be the major determinant of erythrocyte invasion by the parasite.^2-4^ Although *Pf*AMA1 and *Pf*RON2 are highly polymorphic proteins, their interacting residues are fully conserved across *Pf* strains; and thus have been identified as a promising protein-protein interactions drug target.

There are only a handful of inhibitors known to disrupt *Pf*AMA1-*Pf*RON2 interactions.^2-9^ The native ectodomain of the *Pf*RON2 (*Pf*RON2_2021-2059_) is known to inhibit the parasite (most common strain of *Plasmodium falciparum* 3D7) growth efficiently.^2^ Scanlon and co-workers^4^ reported a 29 residue *Pf*RON2 ectodomain mutant (*Pf*RON2_2027-2055_, F2038W, Q2046M) having a half-maximal inhibitory concentration (IC_50_) of 0.16 μM for 3D7 strain. There are few peptide inhibitors reported against *Pf*AMA1 that are not based on *Pf*RON2 sequence.^5-7^ The best among them is a 20-residue peptide (designated as R1 by Harris et al.)^7^ obtained from phage-display screening using a random peptide library that has an IC_50_ (3D7 strain) of ∼4 μM.^2^ R1 is known to be 3D7 strain-specific and the *Pf*AMA1 polymorphism at Tyr^175^ and Ile^225^ has been found to be the major factor for the loss of R1 specificity.^2^ In this work, we demonstrate a unique approach of hybridizing peptide fragments from R1 and *Pf*RON2 to generate an exceedingly potent *Pf*AMA1 inhibitor. The resultant chimeric peptide exhibited two orders of magnitude higher efficacy compared to any of its parent peptides. Also, the peptide chimera was equally effective against most common *P. falciparum* strain 3D7 and a representative strain Dd2 which carries I225N mutation that can impair R1 specificity.

## Results and Discussion

It is evident from the overlay of the two crystal structures (PDB ID 3SRJ and 3SRI) that the R1_1-20_ peptide and *Pf*RON2_2021-2059_ peptide use the same hydrophobic groove to bind to *Pf*AMA1 (**Figure 1A**).^2^ To estimate the extent of interactions, we compared the energetic contribution per residue obtained from molecular mechanics/Poisson–Boltzmann surface area (MM/PBSA) calculations for both the peptides.^10^ Residue-wise energy contribution was calculated by MM/PBSA using the g_mmpbsa tool.^11^ For MM/PBSA calculations, we used *Pf*AMA1 (3D7 strain) bound crystal structures of R1_1-20_ (PDB: 3SRJ) and RON2_2027-2055_ (PDB: 3SRI) and performed short molecular dynamics (MD) simulations (two repeats of 30 ns). Since two molecules of R1 (R1_A1-20_ and R1_B1-20_) are in the crystal structure of *Pf*AMA1-R1 complex, both of them were taken for MD-MM/PBSA calculations. The final 5 ns from each simulation trajectories were then used to determine the residue-wise free energy contributions for both R1 and RON2. Interestingly, the estimated total free-energy contributions from the first ten residues of R1_A1-10_ and RON2_2027-2036_ fragments were -127.03 kJ/mol and -5.18 kJ/mol, respectively (**Figure 1B**). This data indicates that R1_A1-10_ fragment binds to the *Pf*AMA1 groove much tighter than that of the RON2_2027-2036_. On the other hand, the combined free-energy contribution from the rest 14 residues of RON2_2037-2050_, spanning the disulfide containing loop region, in the binding groove was -92.88 kJ/mol that is significantly higher than the combined R1_A11-A18_ and R1_B7-B10_ fragments (−40.25 kJ/mol).

**Figure 1.**
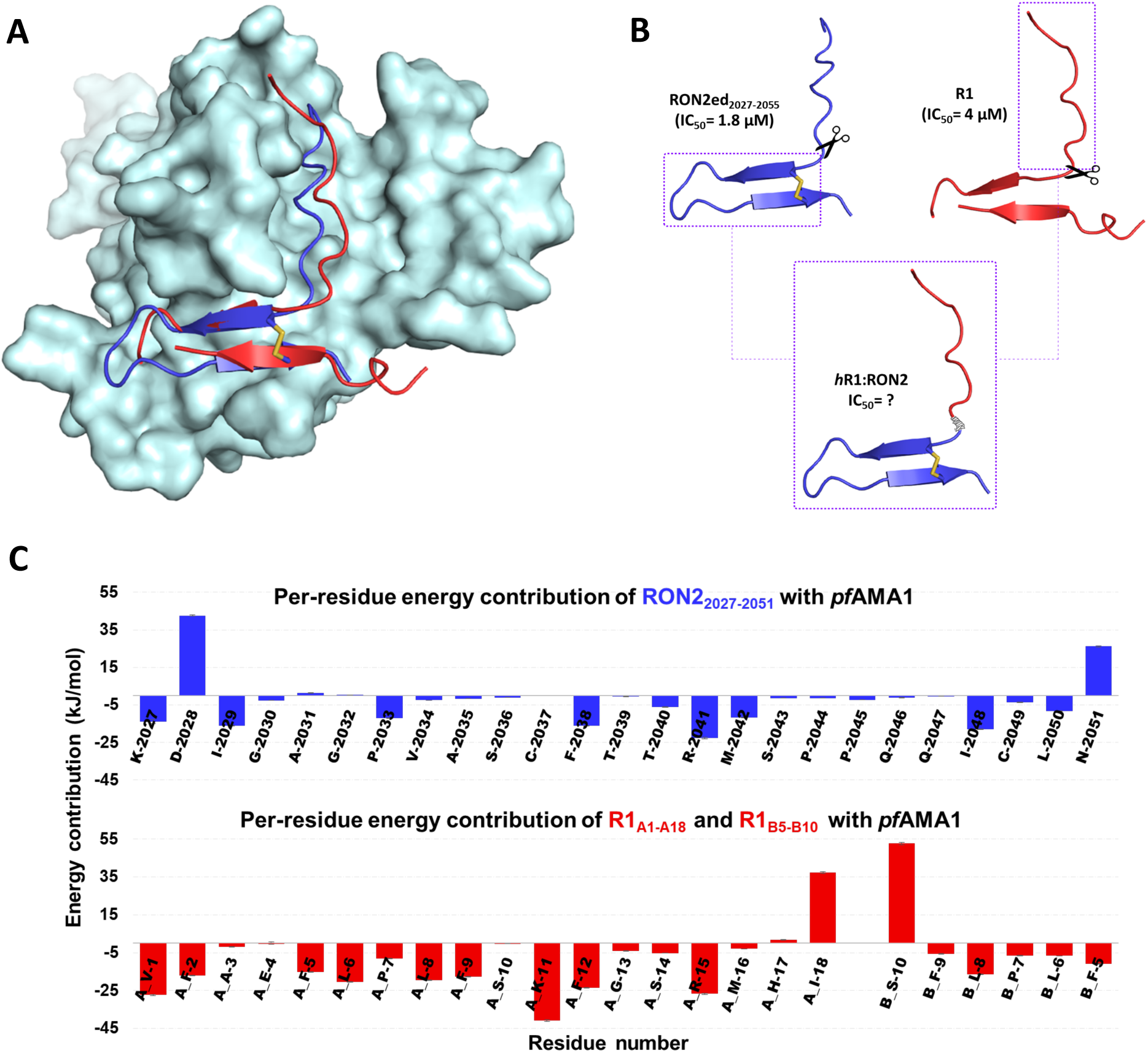
Interactions of R1 and RON2_2027-2055_ with PfAMA1. A) Overlay of the structure of R1 and RON2_2027-2055_ on PfAMA1 groove. B) Schematics showing hybridization of R1 and RON2_2027-2055_. C) The per-residue energy contribution of R1 and RON2_2027-2055_ obtained from MM/PBSA calculations.

Based on the above analysis from crystal structures reported earlier, we designed a small 25-residue chimeric peptide combining the sequences of the high-affinity fragments of both R1 and the peptidic ectodomain of RON2 (**Figure 1C**). The resulting hybrid peptide consisted of the first 10 residues from R1 (R1_1-10_) and all 14 interacting residues from RON2_2037-2050_. Interestingly, the Leu^2050^ of RON2_2027-2055_ occupied the same hydrophobic pocket of *Pf*AMA1 as that of the Leu^B6^ of R1, contributing favourably towards the binding. To avoid unwanted deamidation, in the designed sequence we replaced the native Asn^2051^ of RON2_2027-2055_ by a Ser residue. Consequently, the final designed hybrid peptide (denoted here as R1R2) had total 25 residues. The first 10 residues were from *N*-terminus of the R1 peptide (R1_1-10_) and the next 14 residues were from RON2 (RON2_2037-2050_) with an acetylated *N*-terminus and a free serine residue at the *C*-terminus (Ac-VFAEFLPLFSCFTTRMSPPQQICLS-OH).

Next, we chemically synthesized the R1R2 peptide by machine-assisted Fmoc chemistry solid phase peptide synthesis (SPPS) following a standard protocol at elevated temperature (50°C).^12^ The intra-chain disulfide bond of the hybrid-peptide was formed by air oxidation at pH 9.0 (0.1 M Tris buffer) in presence of 3 M Gu.HCl. The formation of the disulfide bond was confirmed by ESI-MS, and the HPLC purification of the crude folding mixture resulted in the desired peptide in 25.4% yield (see SI Section 2.2 and **Figure 2A**). To compare the parasite growth inhibition efficacy, we also chemically synthesized the parent peptide R1 (VFAEFLPLFSKFGSRMHILK-OH).

**Figure 2.**
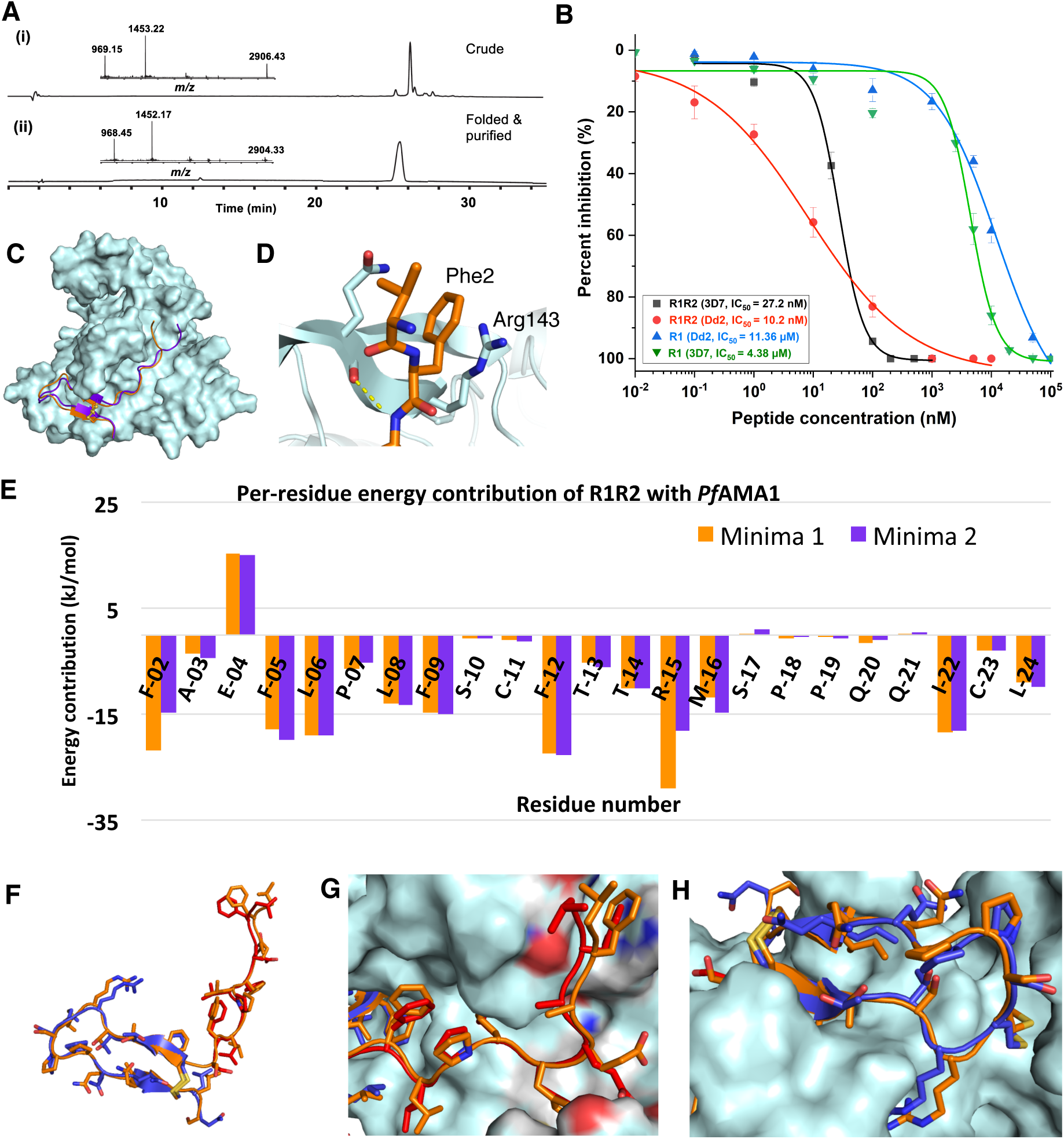
Interactions of hybrid peptide with PfAMA1. A) Analytical RP-HPLC profile (γ = 214 nm) together with ESI-MS data (inset) of R1R2 peptide: (i) unfolded crude; (ii) purified after folding (Observed mass (ESI-MS): 2902.39 ± 0.01 Da (deconvoluted most abundant isotopologue); calculated mass: 2902.40 (most abundant isotopologue)). B) GIA of R1R2 and parent R1 using Pf-3D7 and Pf-Dd2 strains (the data was fitted with nonlinear logistic function for IC50 calculations). C) Superposition of the averaged two energy minimized structures of R1R2-AMA1 for the two minimas (minima 1 (deepest, orange) and minima 2 (purple)). D) Cation-p interaction that drives the lowest energy conformation of R1R2-AMA1 complex. E) The per-residue energy contribution of R1R2 obtained using MM/PBSA for the two minimas reflected in the free energy landscape. The energy contributions from the flexible N- and C-terminal residues (one from each terminus) have not been shown in the plot. F) Superposition of R1R2 (orange) with R1_1-10_ (red) and RON2_2037-2051_ (blue). G) *N*-terminus (first ten residues) of the R1R2 (orange) occupies the same groove of AMA1 as the R11-10 (red). H) *C*-terminus (14 residues) of the R1R2 (orange) occupies the same binding pocket of the AMA1 as the RON2_2037-2051_ (blue).

We then examined the parasite invasion inhibitory activity of the designed hybrid peptide as well as the parent peptides using *ex vivo* growth inhibition assay (GIA) with commonly observed *P. falciparum* strain 3D7 and a representative drug-resistant strain Dd2. The GIAs were performed using *Pf-*3D7 and *Pf-*Dd2 culture with 0.4% and 0.2% parasitemia, respectively, following the protocol described elsewhere (details are given in SI Section 3.1).^2,12-13^

It is important to note that the reported IC_50_^3D7^ of the parent peptides, RON2_2027-2055_ and R1, are 1.8 μM and 4 μM, respectively.^2,4^ Whereas, the designed hybrid peptide R1R2 exhibited an IC_50_^3D7^ of 15.7 nM (SYBR Green fluorescence assay, Figure S5) and 27.2 nM (Giemsa blood smear assay) in the inhibition assay, which is around hundredfold better than its parent peptides. Besides, 25 residue R1R2 exhibited ∼6 fold better IC_50_^3D7^ than the best-known inhibitor against *Pf*AMA1 till date, the 29 residue RON2_2027-2055_ mutant (F2038W, Q2046M; IC_50_^3D7^: 0.16 μM).^4^ Additionally, the hybrid peptide showed an order of magnitude better inhibitory activity than the native 39 residue *Pf*RON2 ectodomain, *Pf*RON2_2021-2059_, which has IC_50_^3D7^ of 151.7 nM (SYBR Green, Figure S5) and >0.2 μM (Giemsa).^2^ Further, to examine the effect, if any, of Scanlon’s mutations^4^ (F2038W, Q2046M) on the invasion inhibition activity, we introduced F12W and Q20M mutations to the designed hybrid peptide (details are given in SI Section 2.3). However, the hybrid peptide with Scanlon’s mutations, R1R2_mut_ (F12W, Q20M), showed 3 times lower inhibitory activity compared to the R1R2 (Figure S5), but still better than other known peptide inhibitors.

Next, we examined the inhibitory activity of R1R2 with a representative drug-resistant strain of *P. falciparum, Pf*-Dd2, and compared the results with the parent R1-peptide. The *Pf-*Dd2 growth inhibition assay with R1-peptide exhibited 2.5 times less inhibitory activity (IC_50_^Dd2^: 11.36 μM, Giemsa) than that of the *Pf-3D7* strain (IC_50_^3D7^: 4.38 μM, Giemsa) (**Figure 2B)**. In stark contrast, R1R2 exhibited an IC_50Dd2_ of 10.2 nM (Giemsa) in the inhibition assay, which is 2.5 times better compared to the *Pf-3D7* strain, and three orders-of-magnitude better *Pf-*Dd2 inhibitory activities than its parent R1-peptide. Interestingly, these results evince that the polymorphism of *Pf*AMA1 at position 225 does not deter the inhibitory activity of the hybrid peptide, unlike its parent R1 peptide that exhibits narrow 3D7 specificity.

To gain structural insights on the AMA1-hybrid peptide interactions, we modelled the hybrid peptide R1R2 on *Pf*AMA1 by taking PDB:3ZWZ^2^ as the starting coordinates. For the MD simulation of the R1R2-AMA1 complex the missing domain-II loop in the crystal structure was reconstructed using charmm-gui webserver (www.charmm-gui.org). MD simulations were performed using GROMACS^14^ v.2019 with CHARMM36^15^ force field. Two repeats of 500 ns MD simulation runs were conducted. Subsequently, principal component analysis was performed on the conformational ensemble obtained from MD simulations of R1R2-AMA1 complex (**Figure S7**).^16-19^ The free-energy landscape constructed from principal component analysis using concatenated trajectories obtained from the MD simulations showed two minimas, the deepest (minima 1) being the tail end of the peptide (spanning val1 to phe5) interacting with the beta-hairpin (Gly132-Leu144) of AMA1. In the other minima (minima 2), the peptide tail was found to be residing more into the cavity distant from the previous one (**Figure 2C**). During the simulation, we observed cation-pi interaction involving Phe2 of the ligand and Arg143 of the AMA1 protein that initiated the formation of intermolecular H-bond along the backbone between Ala3 and Gln141 of the beta-sheet of AMA1 protein, guiding the complex towards the deepest minima (**Figure 2D**).

We then examined the influence of the polymorphic site Tyr^175^ and Ile^225^, the determinant residues for the 3D7 specificity of R1, on the interactions of R1R2 with the *Pf*AMA1 (**Figure 3)**. It is evident from the crystal structure of R1-AMA1 complex (PDB:3SRJ) that the Tyr^175^ makes a hydrogen bond with Leu^6^ backbone of R1, leading to reduced affinity of R1 towards other Tyr^175^ mutant strains.^2^ A careful look at MD trajectories of our R1R2-AMA1 complex revealed that the Tyr^175^ maintained a significant distance from the Leu^6^ of R1R2 throughout, suggesting that the mutation at position 175, as observed in several R1 resistant strains, would not affect the binding of R1R2 with the *Pf*AMA1 (**Figure 3C-D**). It is also important to note here that the mutation at position 225 adversely affects binding specificity of R1 but does not impact the binding of RON2 peptide to *Pf*AMA1 protein.^2^ Since the polymorphic site Ile225 is located near the *Pf*RON2 based fragment of the hybrid peptide R1R2, we do not anticipate any loss of specificity of R1R2 to the parasite strains that carry mutation at position 225. Indeed, we observed three orders-of-magnitude better inhibitory activities against *Pf-*Dd2 (a mutant strain that carries I225N mutation) for the hybrid peptide R1R2 than the parent R1-peptide.

**Figure 3.**
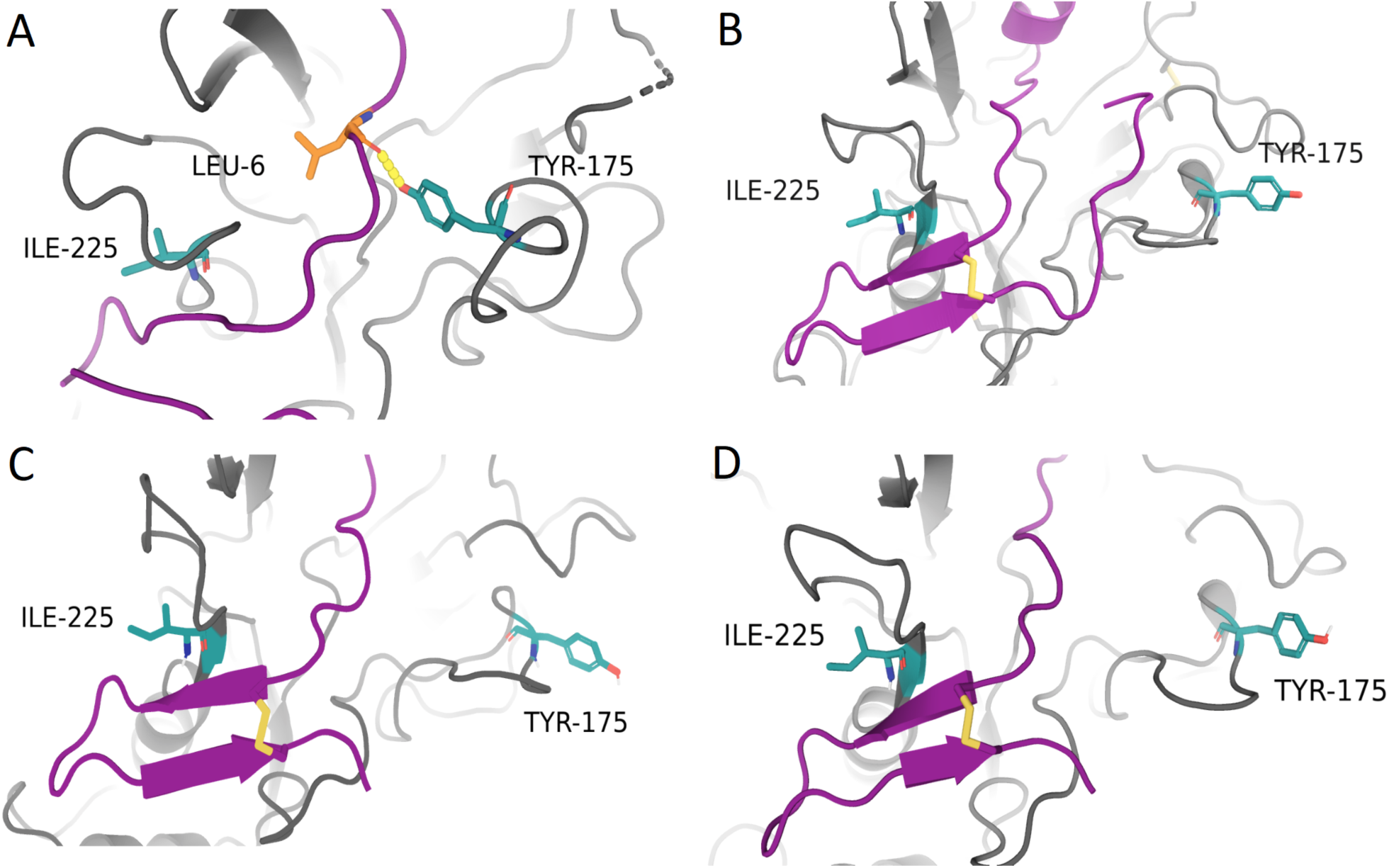
Visualization of polymorphic sites (in yellow) of *Pf*AMA1 (Tyr^175^ and Ile^225^) as observed in the structure of A) R1-AMA1 complex (3SRJ, 3D7), B) RON2-AMA1 complex (3ZWZ, 3D7), C) R1R2-AMA1 complex (MD minima-1, 3D7) and D) R1R2-AMA1 complex (MD minima-2, 3D7).

In order to rationalize the observed augmented potency of the hybrid peptide, we analysed the residue-wise energy contribution of the R1R2 peptide when bound to AMA1. MM/PBSA analysis of the last five-nanosecond simulation trajectories of R1R2-AMA1 complex revealed that the majority of the residues, except Glu4, energetically contributed favorably towards the binding with the AMA1 (**Figure 2E**). Thus, the rational fusion of the best binding features of the two peptides resulted in the extremely high potency of the chimeric peptide that surpassed the potency of any of the parent peptides to a significantly large extent. The superposition of a representative hybrid peptide conformation from the deepest minima with R1_1-10_ and RON2_2037-2051_ showed that the chimeric peptide occupied the same groove of the AMA1 for binding (**Figure 2F-H**).

In summary, the rationally designed peptide chimera, based on R1 and RON2 ectodomain sequence, was found to be two orders-of-magnitude better inhibitor than its parent peptides and an order-of-magnitude better than the native extracellular peptidic domain of the *Pf*RON2. Gratifyingly, the designed high-affinity peptide binder exhibited strong inhibitory activity towards the *Pf-*Dd2 strain, unlike its parent R1-peptide. Further, the hybrid peptide was computationally modeled on the *Pf*AMA1 to understand the molecular basis of its augmented potency. The successful hybridization strategy using the combination of MD-MM/PBSA demonstrated here will open up new possibilities to design novel peptide inhibitors for other PPI based drug targets as well.

## Methods

Experimental methods are described in detail in the Supporting Information.

## Supporting information

Supplementary Informations

## Acknowledgments

We thank late Professor Surajit Sengupta for the critical reading of the initial draft of the manuscript. We are grateful to Dr. Jagannath Mondal for his help on the computational aspects of this work.

## Table of Contents

A peptide chimera displayed malaria parasite growth inhibition with two orders-of-magnitude greater potency than any of its parent peptides

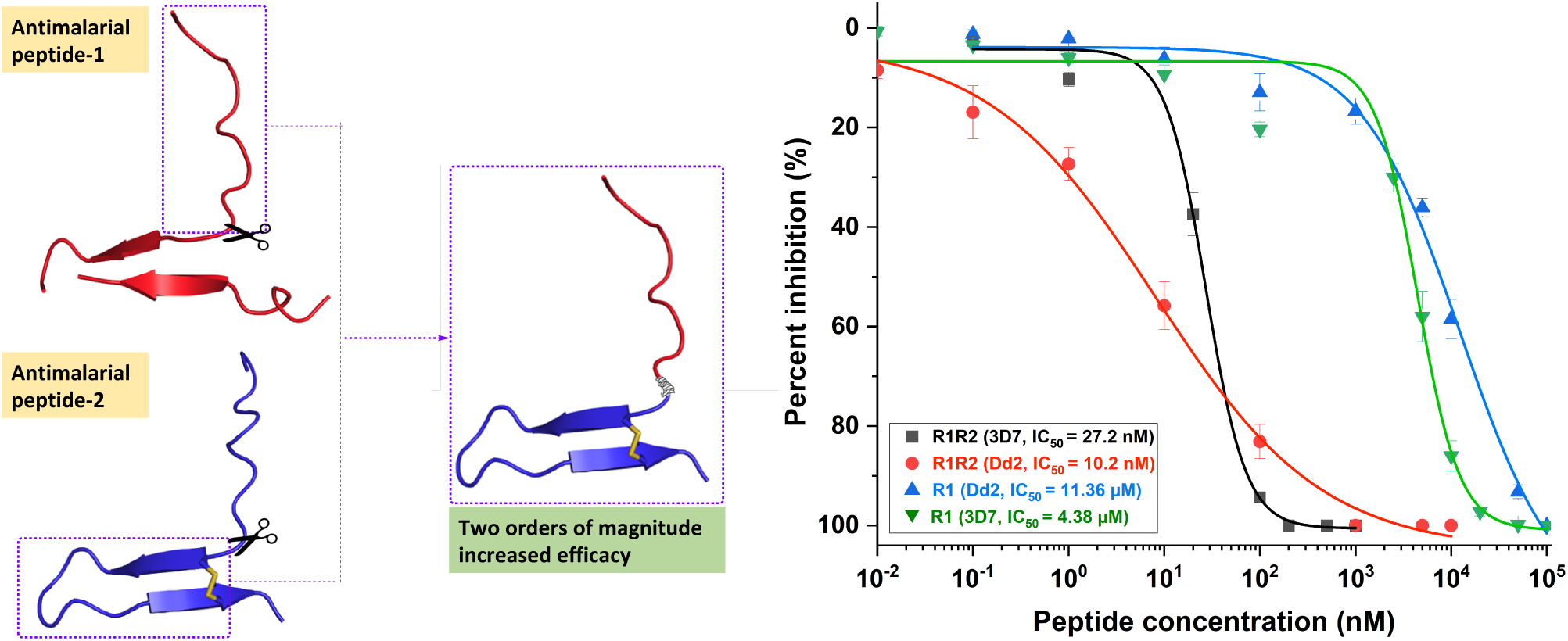

## References

1. World Health, O., World Malaria Report 2018. WHO 2018.

2. Vulliez-Le Normand, B.; Tonkin, M. L.; Lamarque, M. H.; Langer, S.; Hoos, S.; Roques, M.; Saul, F. A.; Faber, B. W.; Bentley, G. A.; Boulanger, M. J.; Lebrun, M., Structural and Functional Insights into the Malaria Parasite Moving Junction Complex. PLoS Pathog. 2012, 8 (6), e1002755.

3. Srinivasan, P.; Yasgar, A.; Luci, D. K.; Beatty, W. L.; Hu, X.; Andersen, J.; Narum, D. L.; Moch, J. K.; Sun, H.; Haynes, J. D.; Maloney, D. J.; Jadhav, A.; Simeonov, A.; Miller, L. H., Disrupting malaria parasite AMA1-RON2 interaction with a small molecule prevents erythrocyte invasion. Nat. Commun. 2013, 31, 2261.

4. Wang, G.; Drinkwater, N.; Drew, D. R.; MacRaild, C. A.; Chalmers, D. K.; Mohanty, B.; Lim, S. S.; Anders, R. F.; Beeson, J. G.; Thompson, P. E.; McGowan, S.; Simpson, J. S.; Norton, R. S.; Scanlon, M. J., Structure–Activity Studies of β-Hairpin Peptide Inhibitors of the Plasmodium falciparum AMA1–RON2 Interaction. Journal of Molecular Biology 2016, 428 (20), 3986–3998.

5. Li, F.; Dluzewski, A.; Coley, A. M.; Thomas, A.; Tilley, L.; Anders, R. F.; Foley, M., Phage-displayed peptides bind to the malarial protein apical membrane antigen-1 and inhibit the merozoite invasion of host erythrocytes. J Biol Chem 2002, 277 (52), 50303–10.

6. Keizer, D. W.; Miles, L. A.; Li, F. M.; Nair, M.; Anders, R. F.; Coley, A. M.; Foley, M.; Norton, R. S., Structures of phage-display peptides that bind to the malarial surface protein, apical membrane antigen 1, and block erythrocyte invasion. Biochemistry-Us 2003, 42 (33), 9915–9923.

7. Harris, K. S.; Casey, J. L.; Coley, A. M.; Masciantonio, R.; Sabo, J. K.; Keizer, D. W.; Lee, E. F.; McMahon, A.; Norton, R. S.; Anders, R. F.; Foley, M., Binding hot spot for invasion inhibitory molecules on Plasmodium falciparum apical membrane antigen 1. Infect Immun 2005, 73 (10), 6981–9.

8. Harris, K. S.; Casey, J. L.; Coley, A. M.; Karas, J. A.; Sabo, J. K.; Tan, Y. Y.; Dolezal, O.; Norton, R. S.; Hughes, A. B.; Scanlon, D.; Foley, M., Rapid Optimization of a Peptide Inhibitor of Malaria Parasite Invasion by Comprehensive N-Methyl Scanning. J Biol Chem 2009, 284 (14), 9361–9371.

9. Akter, M.; Drinkwater, N.; Devine, S. M.; Drew, S. C.; Krishnarjuna, B.; Debono, C. O.; Wang, G.; Scanlon, M. J.; Scammells, P. J.; McGowan, S.; MacRaild, C. A.; Norton, R. S., Identification of the binding site of apical membrane antigen 1 (AMA1) inhibitors using a paramagnetic probe. ChemMedChem 2019, 14 (5), 603–612.

10. Homeyer, N.; Gohlke, H., Free Energy Calculations by the Molecular Mechanics Poisson-Boltzmann Surface Area Method. Mol. Inform. 2012, 31 (2), 114–22.

11. Kumari, R.; Kumar, R.; Open Source Drug Discovery, C.; Lynn, A., g_mmpbsa--a GROMACS tool for high-throughput MM-PBSA calculations. J Chem Inf Model 2014, 54 (7), 1951–62.

12. Mannuthodikayil, J.; Singh, S.; Biswas, A.; Kar, A.; Tabassum, W.; Vydyam, P.; Bhattacharyya, M. K.; Mandal, K., Benzimidazolinone-Free Peptide o-Aminoanilides for Chemical Protein Synthesis. Org Lett 2019, 21 (22), 9040–9044.

13. Vydyam, P.; Dutta, D.; Sutram, N.; Bhattacharyya, S.; Bhattacharyya, M. K., A small-molecule inhibitor of the DNA recombinase Rad51 from Plasmodium falciparum synergizes with the antimalarial drugs artemisinin and chloroquine. J Biol Chem 2019, 294 (20), 8171–8183.

14. Abraham, M. J.; Murtola, T.; Schulz, R.; Páll, S.; Smith, J. C.; Hess, B.; Lindahl, E., GROMACS: High performance molecular simulations through multi-level parallelism from laptops to supercomputers. SoftwareX 2015, 1-2, 19–25.

15. Best, R. B.; Zhu, X.; Shim, J.; Lopes, P. E.; Mittal, J.; Feig, M.; Mackerell, A. D., Jr., Optimization of the additive CHARMM all-atom protein force field targeting improved sampling of the backbone phi, psi and side-chain chi(1) and chi(2) dihedral angles. J. Chem. Theory Comput. 2012, 8 (9), 3257–3273.

16. Kitao, A.; Go, N., Investigating protein dynamics in collective coordinate space. Curr. Opin. Struct. Biol. 1999, 9 (2), 164–9.

17. Amadei, A.; Linssen, A. B.; Berendsen, H. J., Essential dynamics of proteins. Proteins 1993, 17 (4), 412–25.

18. Berendsen, H. J.; Hayward, S., Collective protein dynamics in relation to function. Curr. Opin. Struct. Biol. 2000, 10 (2), 165–9.

19. Kubitzki, M. Enhanced Conformational Sampling of Proteins Using TEE-REX, Chapter 4. University of Goettingen, 2007.

