## Supplementary Informations for "A Chimeric Peptide Inhibits Red Blood Cell Invasion by Plasmodium falciparum with Hundredfold Increased Efficacy"

1. General Methods

### 1.1. Materials and Reagents

Ethyl cyanohydroxyiminoacetate (Oxyma), Guanidine hydrochloride (Gu.HCl), N,N-Diisopropylethylamine (DIEA), all the Nα-Fmoc protected amino acids (the side-chain protecting groups used were, Asp(OtBu), Glu(OtBu), Asn(Trt), Gln(Trt), Lys(Boc), Arg(Pbf), Ser(tBu), and Thr(tBu)) were obtained from Chem-Impex International, USA. Fmoc-L-Cys(trt)-OH was purchased from Gyros Protein Technologies. N,N-Dimethylformamide (DMF), dichloromethane(DCM), diethyl ether, trifluoroacetic acid (TFA) and N,N'-diisopropylcarbodiimide (DIC) were purchased from SRL chemicals India. The HPLC grade acetonitrile (CH3CN) for peptide purification was purchased from Thermofisher Scientific, India. Piperidine was obtained from AVRA chemicals, India. 2-Chlorotrityl chloride (2-Cl-(Trt)-Cl) resin was purchased from Supra Sciences, India. All other common reagents were purchased from Sigma-Aldrich and were of the purest grade available.

### 1.2. Reverse-phase HPLC and LC-MS analysis

Analytical reverse-phase (RP) HPLC was performed on an Agilent HPLC instrument using an Agilent zorbax SB-C3 (5 µm), 4.6×150 mm reverse-phase silica column at a flow rate of 0.9 mL/min using a linear gradient of 10-54% solvent B in solvent A over 22 min or 10-64% solvent B in solvent A over 27 min at 40 °C (solvent A= 0.1% TFA in $H_2O$; solvent B = 0.08% TFA in acetonitrile). The UV absorbance of the column eluant was monitored at 214 nm wavelength. The peptide masses were measured by on-line LC-MS using an Agilent 1290 infinity II/6530 Q-TOF LC/MS instrument. Calculated masses were based on the most abundant isotopologue predicted by Agilent Isotope Distribution Calculator (vr. 7.0.7024.0) software unless stated otherwise. Agilent MassHunter Qualitative Analysis (vr. B.07.00) software was used for the deconvolution mass from the charge states and reported the most abundant isotope observed.
Preparative reverse phase HPLC of crude peptides was performed with a Waters 1525 preparative HPLC system using Waters C4 (5 µm, 300 Å, 10 x 250 mm) or Agilent ZORBAX-SB C3 (5 µm, 80 Å, 9.4 x 250 mm) columns at 40 °C using an appropriate shallow gradient of increasing concentration of solvent B in solvent A at a flow rate of 5 mL/min. Fractions containing the purified target peptide were identified by ESI-MS. Selected pure fractions were then pooled and lyophilized.

### 1.3. A general protocol for machine-assisted Fmoc-SPPS

Fmoc-SPPS was performed by following reported protocol with minor modifications.[1-2] All the peptides were synthesized using an automated peptide synthesizer (Tribute-UV/IR from Protein Technologies, USA). Each steps of the coupling were carried out using amino acids (AA) (0.25 M), DIC (0.25 M) as a coupling reagent and Oxyma (0.25 M) with DIEA (0.025 M) as additives, unless otherwise mentioned. Cysteine and histidine was coupled for 2 min at room temperature followed by 5 min at 50 °C and Arginine was coupled for 20 min at room temperature followed by 5 min at 60 °C. All other amino acid coupling on the 2-Cl-(Trt)-Cl resin was performed for 6 min at 50 °C under $N_2$ atmosphere. Fmoc deprotection after every coupling cycle was carried out by 20% piperidine treatment at 50 °C. After synthesis, the peptides were cleaved from the resin using TFA (85%), Phenol (5%), TIPS (2.5%), Water (5%) and DODT (2.5%) as a cleavage cocktail. After cleavage, the TFA was evaporated under $N_2$ flow inside a well-ventilated fume hood. The cleaved peptide was precipitated and washed with diethyl ether. Dry crude peptides were dissolved in 6 M Gu.HCl and loaded directly on to a preparative HPLC column for purification.

### 2. Chemical synthesis of peptides

#### 2.1. Parent R1-Peptide

The R1-peptide, $NH_2$-Val-Phe-Ala-Glu-Phe-Leu-Pro-Leu-Phe-Ser-Lys-Phe-Gly-Ser-Arg-Met-His-Ile-Leu-Lys-COOH, was synthesized using 2-Cl-(Trt)-Cl resin (scale = 0.125 mmol; substitution = 0.453 mmol/g) by stepwise Fmoc chemistry SPPS in an automated peptide synthesizer at 50 °C (see **Section-1.3** for the peptide synthesis protocol). After global deprotection using the TFA cocktail, the crude peptide was precipitated using diethyl ether. Purification of the crude peptide by preparative HPLC gave 108.6 mg (45 µmol, 37%) of the pure R1-peptide. Observed mass (**ESI-MS**): 2367.30 ± 0.01 Da (deconvoluted most abundant isotopologue); calculated mass: 2367.30 Da (most abundant isotopologue) (**Figure S1**).

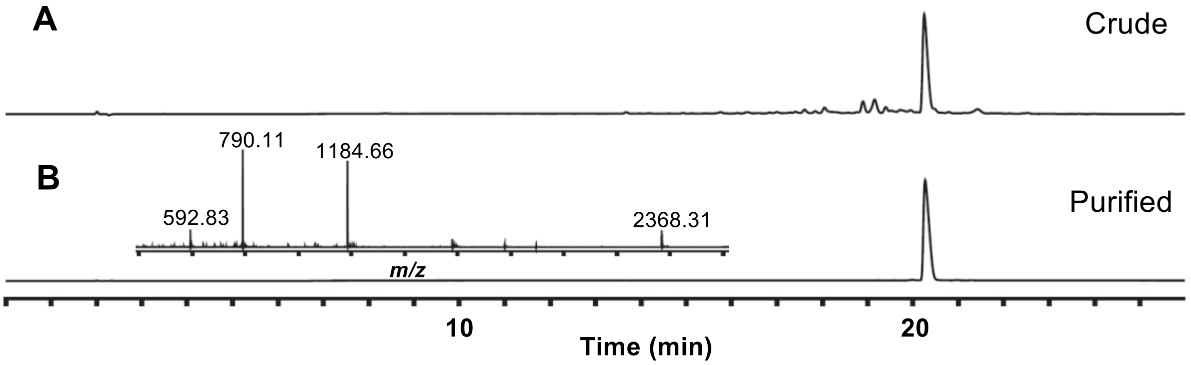

**Figure S1.** Analytical HPLC profile (λ= 214 nm) together with ESI-MS data (inset) of the parent R1-peptide. (A) Crude R1-peptide. (B) Purified peptide R1-peptide. Linear gradient 10%-54% of B over 22 min including 4 min equilibration using an Agilent Zorbax SB-C3, 5 µm, 4.6 x 150 mm, LC column with 0.9 mL/min flow rate was used for the chromatographic separation. Purification was performed using a linear gradient 22%-42% of buffer B in buffer A over 80 min with a flow rate of 5 mL/min at 40 °C (buffer A = 0.1% acetic acid in water; buffer B = 0.08% acetic acid in acetonitrile) using a C12, 10 x 250 mm column (Phenomenex-Proteo, 90 Å, 5 µm).

#### 2.2. Designed Hybrid Peptide, R1R2

The designed hybrid peptide, Ac-Val-Phe-Ala-Glu-Phz-Leu-Pro-Leu-Phe-Ser-Cys-Phe-Thr-Thr-Arg-Met-Ser-Pro-Pro-Gln-Gln-Ile-Cys-Leu-Ser-COOH, was synthesized using 2-Cl-(Trt)-Cl resin (scale = 0.3 mmol; substitution = 0.453 mmol/g) by stepwise Fmoc chemistry SPPS in an automated peptide synthesizer at 50 °C (see **Section-1.3** for the peptide synthesis protocol). After global deprotection using the TFA cocktail, the crude R1R2 was precipitated using diethyl ether. Observed mass (**ESI-MS**): 2904.42 ± 0.01 Da (deconvoluted most abundant isotopologue); calculated mass: 2904.41 Da (most abundant isotopologue) (**Figure S2A**). For oxidative folding, 0.15 mmol (~370 mg) of the crude R1R2 was dissolved in 100 mL of a folding buffer consisting of 0.1 M Tris (pH 8.9), 3 M Gu.HCl and incubated at room temperature for 48 Hrs. Progress of the folding reaction was monitored by LCMS. After the completion of the folding, the reaction mixture was acidified to pH 3 by 6 M HCl and purified through HPLC yielding 110.9 mg (38.2 µmol, 25.4% yield) of the desired folded R1R2**.** Observed mass **(ESI-MS)**: 2902.39 ± 0.01 Da (deconvoluted most abundant isotopologue), calculated mass: 2902.40 Da (most abundant isotopologue) **(Figure S2B).**

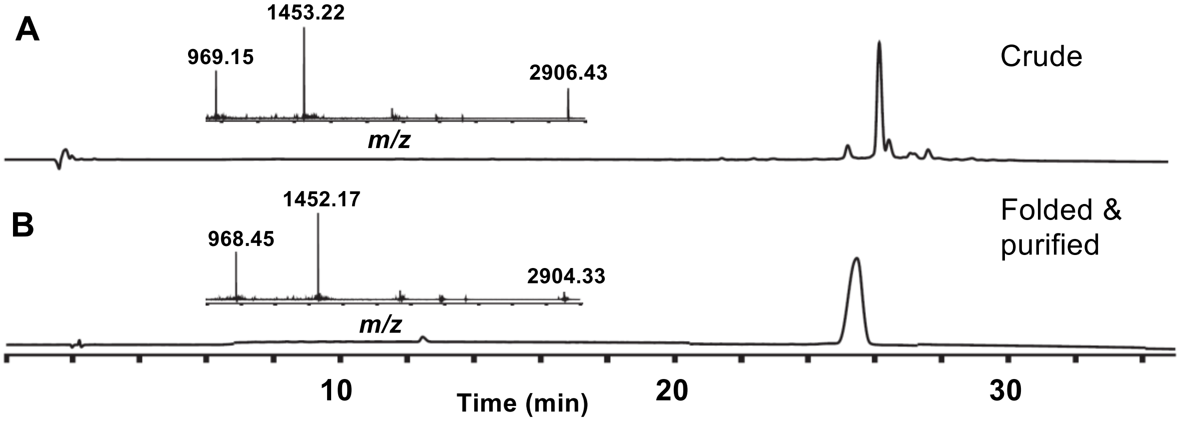

**Figure S2.** Analytical HPLC profile (λ= 214 nm) together with ESI-MS data (inset) of the designed hybrid peptide. (A) Crude R1R2. (B) Folded and purified peptide R1R2. Linear gradient 10%-70% of B over 30 min including 4 min equilibration using Agilent Zorbax SB-C3, 5 µm, 4.6 x 150 mm, LC column with 0.9 mL/min flow rate was used for the chromatographic separation. Purification was performed using a linear gradient 40%-60% of buffer B in buffer A over 40 min with a flow rate of 5 mL/min at 40 °C (buffer A = 0.1% acetic acid in water; buffer B = 0.08% acetic acid in acetonitrile) using a C12, 10 x 250 mm column (Phenomenex-Jupiter-Proteo, 90 Å, 5 µm).

### 2.3. Mutant Hybrid Peptide, R1R2$_{\mathbf{mut}}$ (F12W, Q20M)

The hybrid peptide with F12W, Q20M mutations, Ac-Val-Phe-Ala-Glu-Phz-Leu-Pro-Leu-Phe-Ser-Cys-Trp-Thr-Thr-Arg-Met-Ser-Pro-Pro-Met-Gln-Ile-Cys-Leu-Ser-COOH, was synthesized using 2-Cl-(Trt)-Cl resin (scale = 0.3 mmol; substitution = 0.453 mmol/g) by stepwise Fmoc chemistry SPPS in an automated peptide synthesizer at 50 °C (see **Section-1.3** for the peptide synthesis protocol). After global deprotection using the TFA cocktail, the crude peptide, R1R2$_{mut}$ (F12W, Q20M), was precipitated using diethyl ether. Observed mass (**ESI-MS**): 2947.41 ± 0.01 Da (deconvoluted most abundant isotopologue); calculated mass: 2947.41 Da (most abundant isotopologue) (**Figure S3A**). For oxidative folding, 0.15 mmol (~100 mg) of the crude R1R2$_{mut}$ was dissolved in 50 mL of folding buffer consisting of 0.1 M Tris (pH 8.4), 3 M Gu.HCl and incubated at room temperature for 48 Hrs. Progress of the folding reaction was monitored by LCMS. After the completion of the folding, the reaction mixture was acidified to pH 3 by 6 M HCl and purified through HPLC to furnish 23.4 mg (8 µmol, 5.2% yield) of the desired folded R1R2$_{mut}$ (F12W, Q20M) peptide**.** Observed mass **(ESI-MS)**: 2945.39 ± 0.01 Da (deconvoluted most abundant isotopologue), calculated mass: 2945.40 Da (most abundant isotopologue) **(Figure S3B).**

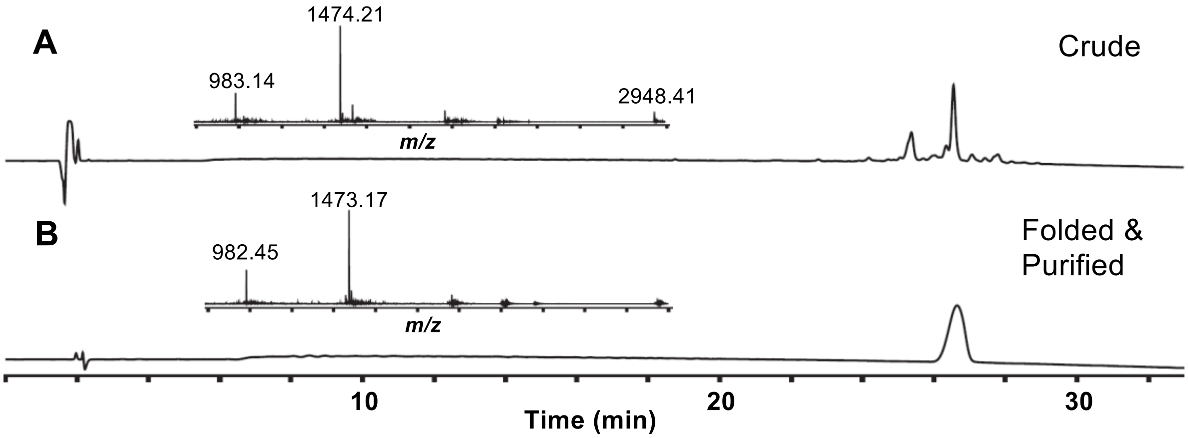

**Figure S3.** Analytical HPLC profile (λ= 214 nm) together with ESI-MS data (inset) of the mutant hybrid peptide. (A) Crude R1R2*mut*. (B) Folded and purified peptide R1R2mut. Linear gradient 10%-70% of B over 30 min including 4 min equilibration using Agilent Zorbax SB-C3, 5 µm, 4.6 x 150 mm, LC column with 0.9 mL/min flow rate was used for the chromatographic separation. Purification was performed using a linear gradient 35%-60% of buffer B in buffer A over 40 min with a flow rate of 5 mL/min at 40 °C (buffer A = 0.1% acetic acid in water; buffer B = 0.08% acetic acid in acetonitrile) using a C4, 10 x 250 mm column (Phenomenex-Jupiter, 300 Å, 10 µm).

### 2.4. The ectodomain of RON2, *Pf*RON2$_{2021\text{-}2059}$

Chemical synthesis, folding, and purification of the ectodomain of RON2, *Pf*RON2$_{2021\text{-}2059}$ has been shown elsewhere (unpublished).

Sequence: $NH_2$-Asp-Ile-Thr-Gln-Gln-Ala-Lys-Asp-Ile-Gly-Ala-Gly-Pro-Val-Ala-Ser-Cys-Phe-Thr-Thr-Arg-Met-Ser-Pro-Pro-Gln-Gln-Ile-Cys-Leu-Asn-Ser-Val-Val-Asn-Thr-Ala-Leu-Ser-COOH

Observed mass **(ESI-MS)**: 4061.02 ± 0.01 Da (deconvoluted most abundant isotopologue), calculated mass: 4060.99 Da (most abundant isotopologue)

### 2.5. Truncated PfRON2 (PfRON2$_{trunc}$)

It is noteworthy to mention that the individual peptide fragments of the hybrid peptide, R1R2$_{1\text{-}11}$ (i.e. R1$_{1\text{-}11}$) and R1R2$_{11\text{-}24}$ (i.e. RON2$_{2037\text{-}2050}$) were previously shown to have no significant growth inhibitory activity.[3-4] However, a 24 residue truncated native RON2 sequence (RON2$_{2027\text{-}2050}$) with acetylated *N*-terminus and a free Serine residue at the *C*-terminus gave 2-fold increase in IC50[3d7] (0.85 µM) compared to the reported RON2$_{2027\text{-}2055}$ (Figure S6).

The truncated *Pf*RON2 (*Pf*RON2$_{trunc}$), Ac-Lys-Asp-Ile-Gly-Ala-Gly-Pro-Val-Ala-Ser-Cys-Phe-Thr-Thr-Arg-Met-Ser-Pro-Pro-Gln-Gln-Ile-Cys-Leu-Ser-COOH, was synthesized using 2-Cl-(Trt)-Cl resin (scale = 0.15 mmol; substitution = 0.453 mmol/g) by stepwise Fmoc chemistry SPPS in an automated peptide synthesizer at 50 °C (see **Section-1.3** for the peptide synthesis protocol). After global deprotection using the TFA cocktail, the crude *Pf*RON2$_{trunc}$, was precipitated using diethyl ether. Observed mass (**ESI-MS**): 2649.27 ± 0.01 Da (deconvoluted most abundant isotopologue); calculated mass: 2649.28 Da (most abundant isotopologue) (**Figure S4A**). For oxidative folding, 0.15 mmol (237 mg) of the crude *Pf*RON2$_{trunc}$ was dissolved in 100 mL of folding buffer consisting of 0.1 M Tris (pH 9.5), 2 M Gu.HCl and incubated at room temperature for 48 Hrs. Progress of the folding reaction was monitored by LCMS. After the completion of the folding, the reaction mixture was acidified to pH 3 by 6 M HCl and purified through HPLC to give 85.6 mg (32.3 µmol, 21.5% yield) of the desired folded *Pf*RON2$_{trunc}$. Observed mass **(ESI-MS)**: 2647.29 ± 0.01 Da (deconvoluted most abundant isotopologue), calculated mass: 2647.26 Da (most abundant isotopologue) **(Figure S4B).**

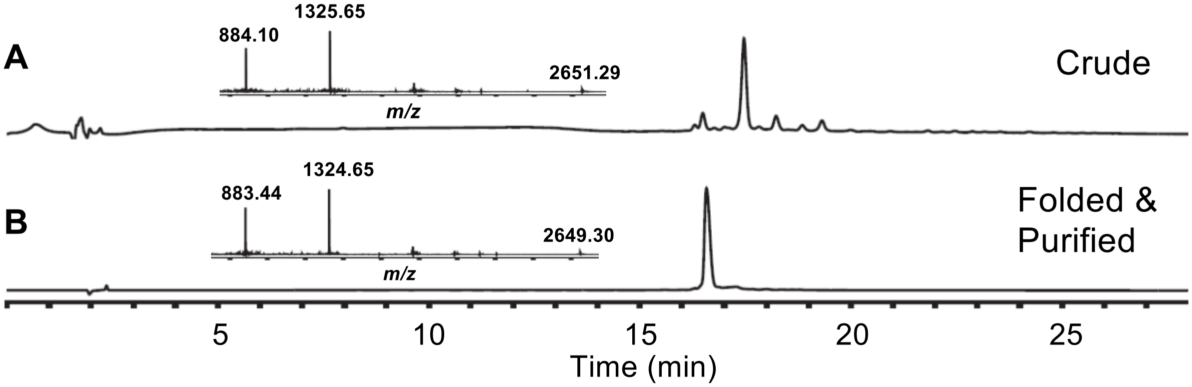

**Figure S4.** Analytical HPLC profile (λ= 214 nm) together with ESI-MS data (inset) of the Truncated *Pf*RON2. (A) Crude *Pf*RON2$_{trunc}$. (B) Folded and purified peptide *Pf*RON2$_{trunc}$. Linear gradient 10%-70% of B over 30 min including 4 min equilibration using Agilent Zorbax SB-C3, 5 µm, 4.6 x 150 mm, LC column with 0.9 mL/min flow rate was used for the chromatographic separation. Purification was performed using a linear gradient 01%-41% of buffer B in buffer A over 40 min with a flow rate of 5 mL/min at 40 °C (buffer A = 0.1% acetic acid in water; buffer B = 0.08% acetic acid in acetonitrile) using a C12, 10 x 250 mm column (Phenomenex-Jupiter-Proteo, 90 Å, 5 µm).

### 3. *Plasmodium falciparum* (*Pf*) Growth inhibition assay (GIA)

#### 3.1. General protocol[2, 5]

Growth Inhibition Assays were performed using reported protocol with minor modifications, as described elsewhere.[2, 5] *Plasmodium falciparum* strains (3D7 & Dd2) were cultured using the candle jar method in RPMI-1640 media supplemented with 1% albumax and 0.005% hypoxanthine. The hematocrit of RBC was 5%. Tightly synchronous (90-95%) culture at the trophozite stage having ~0.2-0.4% of parasitemia was used for all the experiments. Peptide stocks were prepared in DMSO. Cells were treated with increasing concentrations of peptides (R1R2 = 0.001 nM to 10 µM, R1R2$_{mut}$ = 0.001 nM to 10 µM, R1 = 0.01 nM to 100 µM, *PfRON2*$_{2021\text{-}2059}$ = 0.001 nM to 10 µM, and *Pf*RON2$_{trunc}$ = 0.01 nM to 100 µM) and incubated for 72 hours. DMSO was used as vehicle control throughout the experiment. Either Giemsa staining based method and/or SYBR green method was used to measure the percentage parasitemia.

**Giemsa blood smear method**

A thin blood smear was prepared and stained with Giemsa stain after methanol fixation. The infected erythrocytes were then counted in random microscopic fields to evaluate the percentage of parasitemia.

**SYBR green method**

The SYBR green-I based parasite quantification was performed by taking 100 µl of the respective peptide treated parasite culture from each experimental well into a 96 well plate. To this, an equal volume of lysis buffer having

SYBR green-1 (20 mM Tris (pH 7.5), 5 mM EDTA, 0.008% saponin, 0.08% Triton X-100 and SYBR green (1:5000 dilution) were added and incubated at room temperature in dark for 1 hour. Fluorescence intensities were measured with the multimode reader (Bio-Tek Synergy Mx microplate) at excitation and emission wavelengths of 485 nm and 530 nm, respectively. The background reading for the parasite-free medium was subtracted to obtain the correct fluorescence value. Triplicate wells for each dilution of respective peptide were measured, and from these values, the percentage of parasite inhibition was calculated and plotted against the respective peptide concentration.

#### 3.2 Growth inhibition assay graphs

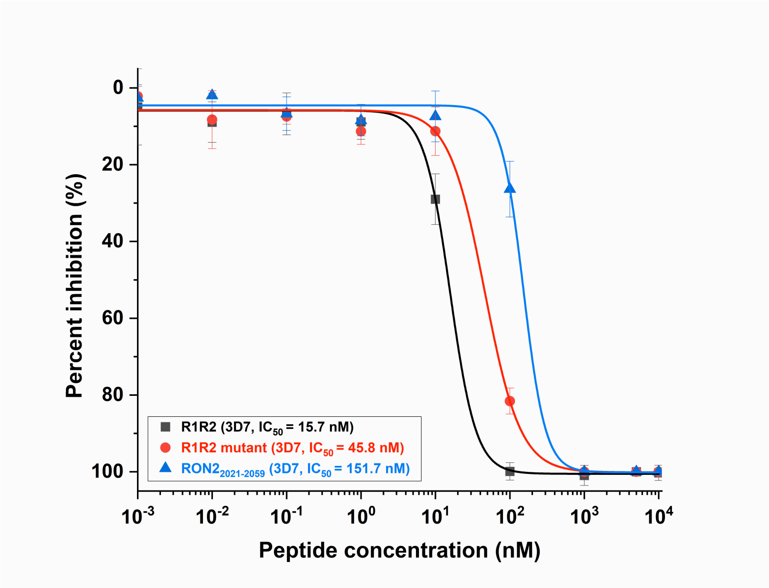

**Figure S5.** Growth inhibition assay (SYBR green method) of R1R2, R1R2$_{mut}$ and *PfRON2$_{2021-2059}$* using *Pf*3D7 strains. The data was fitted with nonlinear logistic function for $IC_{50}$ calculations.

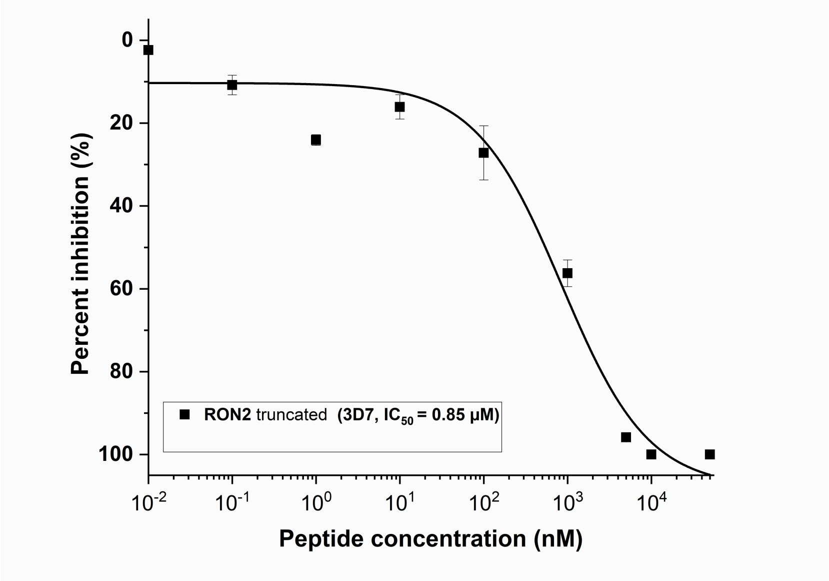

**Figure S6.** Growth inhibition assay (Giemsa blood smear method) of RON2$_{trunc}$ using *Pf*3D7 strains. The data was fitted with nonlinear logistic function for $IC_{50}$ calculations.

### 4. Computational Chemistry

#### 4.1. Molecular Dynamics Simulations

The molecular dynamics simulations were performed using the GROMACS code version 2019[6] with CHARMM36 force field[7] and TIP3P water model. Charmm gui web-server[8] was used to prepare the systems including the construction of the loop. Coordinates from PDB:3ZWZ[9] were used as a starting structure for simulations. A cubic box type with a distance of 1.0 nm between the complex and the box edge was defined to set the minimum distance of at least 2 nm between any two periodic images of the solute. Then, TIP3P[10] water model filled the box and the system was neutralized with potassium chloride counter ions in a concentration of 0.15 mol/L. The steepest descent algorithm was applied to optimize the system during a 100 ps run time. A position restraint of 1000 kJ $mol^{-1}$ $nm^{-2}$ was applied to fix the positions of both ligand and protein atoms and the whole system was equilibrated under the NVT ensemble through 100 ps time during which the V-rescale[11] thermostat was used to regulate the temperature close to 310 K. Following the NVT step, the system was equilibrated under the NPT ensemble through 100-ps time in such a way that the pressure of the system was stabilized to 1 atm. The production MD of 500 ns (two repeats) was performed on the well-equilibrated system at the temperature fixed at 310 K and the pressure of 1 atm. The Particle-mesh Ewald (PME)[12] algorithm calculated the long-ranged electrostatic contributions and the length of all covalent bonds was constrained using the LINCS[13] algorithm, a faster algorithm in comparison to the SHAKE algorithm. Upon termination of the simulation, the protein was backed and cantered in the box along with removing of periodic boundary condition from the trajectory. Subsequently, principal component analysis (PCA) was performed on the conformational ensemble obtained from MD simulations of R1R2-AMA1 complex (**Figure S7**). PCA was constructed for the displacement of Cα atoms based on the covariance matrix.[14-17]). The conformational changes in protein were derived from free energy landscape (FEL) obtained using Pyemma package (http://www.emma-project.org/).

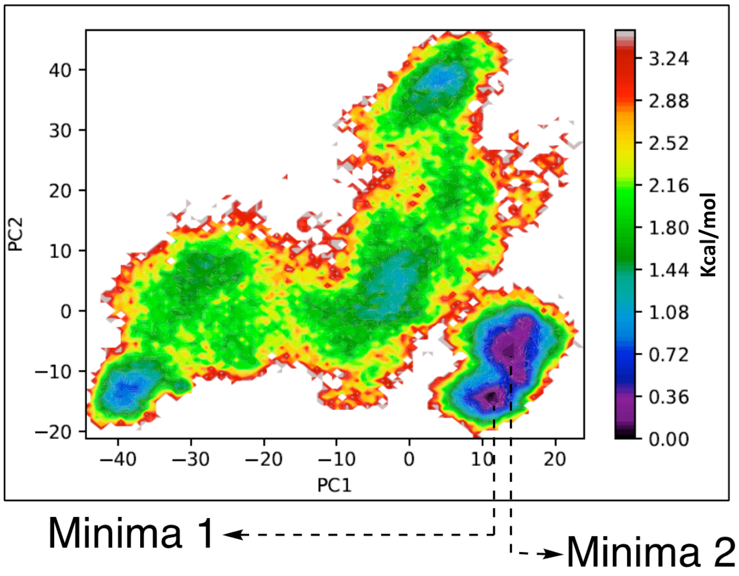

**Figure S7.** FEL was plotted by projecting energy landscapes using the first two principal components, that is, PC1 and PC2, for R1R2-AMA1 complex.

### 4.2. MMPBSA Calculations

The binding free energy of the interactions in the ligand-protein complexes were obtained by utilizing the Molecular Mechanic/Poisson-Boltzmann Surface Area (MMPBSA) method which employs ensembles derived from molecular dynamics (MD) simulation.[18] The g_mmpbsa program[18-19] was used to calculate different components of the binding free energy between AMA1 and all peptide ligands. The last 5 ns of the simulation trajectories were considered for MMPBSA calculations. The free energy of binding was described by the equation; $\Delta G_{binding} = G_{complex} - (G_{protein} + G_{ligand})$, where, $G_{complex}$ is the total free energy of the protein-ligand complex and $G_{protein}$ and $G_{ligand}$ are total free energies of the isolated protein and ligand in solvent, respectively. The free energies for $G_{complex}$, $G_{protein}$, and $G_{ligand}$ were estimated by the equation: $G_x = E_{MM} + G_{solvation}$; where x is the protein, ligand, or complex. $E_{MM}$ is the average molecular mechanics potential energy in vacuum and $G_{solvation}$ is free energy of solvation. The molecular mechanics potential energy was calculated in vacuum as follows:

$E_{MM} = E_{bonded} + E_{non\text{-}bonded}$

$E_{non\text{-}bonded} = E_{vdw} + E_{elec}$

where $E_{bonded}$ is bonded interactions which involves bond, angle, dihedral, and $E_{non\text{-}bonded}$ is non-bonded interactions consisting of van der Waals ($E_{vdw}$) and electrostatic ($E_{elec}$) interactions. $\Delta E_{bonded}$ is always considered as zero. [18] The solvation free energy ($G_{solvation}$) was estimated as the sum of electrostatic solvation free energy ($G_{polar}$) and apolar solvation free energy ($G_{non\text{-}polar}$): $G_{solvation} = G_{polar} + G_{non\text{-}polar}$; where $G_{polar}$ was computed using the Poisson-Boltzmann (PB) equation[18] and $G_{non\text{-}polar}$ was estimated from the solvent-accessible surface area (SASA) using the equation; $G_{non\text{-}polar} = \gamma SASA + b$; where $\gamma$ is a coefficient related to surface tension of the solvent and b is the fitting parameter. The values of the constants are $\gamma = 0.02267$ Kj/mol/Å$^2$ or 0.0054 Kcal/mol/Å$^2$ and b = 3.849 Kj/mol or 0.916 Kcal/mol.

Using the *g_mmpbsa* tool, the binding energy can be decomposed on a per residue basis. Initially the energy components $E_{MM}$, $G_{polar}$, and $G_{nonpolar}$ of individual atoms are calculated in the bound as well as the unbound form, and subsequently their contribution to the binding energy $\Delta R_x^{BE}$ of residue *x* is calculated as follows:

$$\Delta R_x^{BE} = \sum_{i=0}^{n} (A_i^{bound} - A_i^{free})$$

where $A_i^{bound}$ and $A_i^{free}$ are the energy of *i*th atom from *x* residue in bound and unbound forms respectively, and n is the total number of atoms in the residue.

### REFERENCES

1. Collins, J. M.; Singh, S. K. Coupling method for peptide synthesis at elevated temperatures. 2016.

2. Mannuthodikayil, J.; Singh, S.; Biswas, A.; Kar, A.; Tabassum, W.; Vydyam, P.; Bhattacharyya, M. K.; Mandal, K., Benzimidazolinone-Free Peptide o-Aminoanilides for Chemical Protein Synthesis. *Org Lett* **2019**, *21* (22), 9040-9044.

3. Wang, G.; Drinkwater, N.; Drew, D. R.; MacRaild, C. A.; Chalmers, D. K.; Mohanty, B.; Lim, S. S.; Anders, R. F.; Beeson, J. G.; Thompson, P. E.; McGowan, S.; Simpson, J. S.; Norton, R.

S.; Scanlon, M. J., Structure–Activity Studies of β-Hairpin Peptide Inhibitors of the Plasmodium falciparum AMA1–RON2 Interaction. *Journal of Molecular Biology* **2016,** *428* (20), 3986-3998.
4. Wang, G. Q.; MacRaild, C. A.; Mohanty, B.; Mobli, M.; Cowieson, N. P.; Anders, R. F.; Simpson, J. S.; McGowan, S.; Norton, R. S.; Scanlon, M. J., Molecular Insights into the Interaction between Plasmodium falciparum Apical Membrane Antigen 1 and an Invasion-Inhibitory Peptide. *PLoS One* **2014,** *9* (10).
5. Vydyam, P.; Dutta, D.; Sutram, N.; Bhattacharyya, S.; Bhattacharyya, M. K., A small-molecule inhibitor of the DNA recombinase Rad51 from Plasmodium falciparum synergizes with the antimalarial drugs artemisinin and chloroquine. *J Biol Chem* **2019,** *294* (20), 8171-8183.
6. M.J. Abraham; D. van der Spoel; E. Lindahl; B. Hess, *and the GROMACS development team, GROMACS User Manual version 2019,* http://www.gromacs.org.
7. Best, R. B.; Zhu, X.; Shim, J.; Lopes, P. E.; Mittal, J.; Feig, M.; Mackerell, A. D., Jr., Optimization of the additive CHARMM all-atom protein force field targeting improved sampling of the backbone phi, psi and side-chain chi(1) and chi(2) dihedral angles. *J. Chem. Theory Comput.* **2012,** *8* (9), 3257-3273.
8. Jo, S.; Kim, T.; Iyer, V. G.; Im, W., CHARMM-GUI: a web-based graphical user interface for CHARMM. *J. Comput. Chem.* **2008,** *29* (11), 1859-65.
9. Vulliez-Le Normand, B.; Tonkin, M. L.; Lamarque, M. H.; Langer, S.; Hoos, S.; Roques, M.; Saul, F. A.; Faber, B. W.; Bentley, G. A.; Boulanger, M. J.; Lebrun, M., Structural and Functional Insights into the Malaria Parasite Moving Junction Complex. *PLoS Pathog.* **2012,** *8* (6).
10. Jorgensen, W. L.; Chandrasekhar, J.; Madura, J. D.; Impey, R. W.; Klein, M. L., Comparison of Simple Potential Functions for Simulating Liquid Water. *J. Chem. Phys.* **1983,** *79* (2), 926-935.
11. Bussi, G.; Donadio, D.; Parrinello, M., Canonical sampling through velocity rescaling. *J. Chem. Phys.* **2007,** *126* (1).
12. Darden, T.; York, D.; Pedersen, L., Particle Mesh Ewald - an N.Log(N) Method for Ewald Sums in Large Systems. *J. Chem. Phys.* **1993,** *98* (12), 10089-10092.
13. Hess, B.; Bekker, H.; Berendsen, H. J. C.; Fraaije, J. G. E. M., LINCS: A linear constraint solver for molecular simulations. *J. Comput. Chem.* **1997,** *18* (12), 1463-1472.
14. Kitao, A.; Go, N., Investigating protein dynamics in collective coordinate space. *Curr. Opin. Struct. Biol.* **1999,** *9* (2), 164-9.
15. Amadei, A.; Linssen, A. B.; Berendsen, H. J., Essential dynamics of proteins. *Proteins* **1993,** *17* (4), 412-25.
16. Berendsen, H. J.; Hayward, S., Collective protein dynamics in relation to function. *Curr. Opin. Struct. Biol.* **2000,** *10* (2), 165-9.
17. Kubitzki, M. Enhanced Conformational Sampling of Proteins Using TEE-REX, Chapter 4. University of Goettingen, 2007.
18. Kollman, P. A.; Massova, I.; Reyes, C.; Kuhn, B.; Huo, S.; Chong, L.; Lee, M.; Lee, T.; Duan, Y.; Wang, W.; Donini, O.; Cieplak, P.; Srinivasan, J.; Case, D. A.; Cheatham, T. E., 3rd, Calculating structures and free energies of complex molecules: combining molecular mechanics and continuum models. *Acc. Chem. Res.* **2000,** *33* (12), 889-97.
19. Kumari, R.; Kumar, R.; Open Source Drug Discovery, C.; Lynn, A., g_mmpbsa--a GROMACS tool for high-throughput MM-PBSA calculations. *J Chem Inf Model* **2014,** *54* (7), 1951-62.
